# Early Skill Learning Is Shaped by the Offline Emergence of Expert Synergies

**DOI:** 10.64898/2026.02.24.707000

**Authors:** William Kistler, Rawan Fakhreddine, Giselle R Rodriguez, Margaret Hayward, Ethan R. Buch, Sven Bestmann, Leonardo G. Cohen

## Abstract

Everyday skilled actions depend on the formation of coordinated motor synergies that integrate multiple digits into stable, low-dimensional control units. Although initial practice of a new skill leads to rapid performance improvements, it is unclear whether the underlying movement kinematics reorganize on a similar timescale and in a way that directly relates to these gains. It also remains uncertain whether such reorganization occurs mainly during active practice or instead during brief rest breaks. Here, we tracked the temporal evolution of multi-digit synergy formation during early learning of a naturalistic keypress skill. Initial practice rapidly sculpted the motor repertoire toward higher-order, temporally compressed and overlapping multi-digit synergies. These synergies emerged after only minutes of practice and continued to be expressed along the full training session. Notably, they were primarily shaped across brief rest breaks and robustly predicted individual skill proficiency. Across learning, distinct synergy subtypes emerged, differing in their heuristic prevalence. Rarely expressed synergies reflected transient novice patterns, synergies expressed at intermediate levels could index exploratory and trial-initiation strategies, and highly expressed synergies emerged later to dominate performance, reflecting the consolidation and expansion of skilled motor control. Together, these findings indicate that skilled performance is supported by the early formation of a compact repertoire of expert multi-digit synergies that emerge preferentially across rest periods and predict subsequent skill gains. They further raise the hypothesis that explicitly training such expert synergies alongside task goals could enhance learning in domains such as the arts, sport, and neurorehabilitation.

**Highlights:**

Expert multi-digit synergies form during early skill learning
Higher-order synergies emerge across brief rest breaks
Fatigue spares expert synergy use during early learning
Skill gains mirror multi-digit synergy expression

**eTOC blurb:** “Expert multi-digit synergies emerge early in motor learning, strengthen across brief rest breaks, track skill gains, and resist fatigue”

## Introduction

Activities of daily living, from walking and handwriting to athletic play, depend on the precise orchestration of temporally ordered motor sequences ^1–4^. Fluency develops as the motor system learns to coordinate fingers in new ways. These task-specific movement patterns link motions within and across digits so that they work together to meet the demands of the object, the required force, timing, and accuracy ^5,6^. With practice, discrete keypresses become coarticulated into coordinated multi-digit synergies, reducing effective control dimensionality and stabilizing execution, thereby optimizing performance across skills such as grasping ^7–9^, typing ^10^, finger spelling^11^, and haptic exploration ^12^. Studying how motor synergies emerge and reorganize provides a direct window into the neuromotor mechanisms that support the acquisition, refinement, and generalization of skilled actions^13^.

Early learning of naturalistic skills is characterized by prominent initial performance improvements^14–16^. Over the course of learning, skill performance fluctuates during practice and brief rest periods as a result of the dynamic interplay between performance-enhancing processes, such as learning, and performance-limiting mechanisms, including motor or cognitive fatigue (sometimes referred to collectively as reactive inhibition)^17,18^. However, how these competing processes are expressed in the kinematic signatures that drive the transition from novice movement patterns to expert multi-digit synergies, particularly during the earliest phases of skill acquisition, is not known. Early gains in skill may arise from several, non-mutually exclusive, kinematic adjustments: faster execution^19^, greater temporal overlap among digit movements (coarticulation) ^20–22^, reparameterization of velocity profiles^23^, improved spatial accuracy^24^ or their combination. In the temporal domain, the formation of multi-digit synergies characteristic of expertise could unfold rapidly, on the order of seconds, or accrue more slowly over minutes to hours ^25,26^. Such reorganization may also proceed during brief rest intervals^27^ or instead manifest primarily during active practice, in parallel with expansion of memory capacity^28^. This pattern could suggest a dissociation between the time course of behavioral improvement and the optimization of underlying kinematics. In addition, learning processes such as exploration, prospective planning^29^, and expansion of memory capacity^28^ may be distinctly expressed in the transition from novice patterns to expert multi-digit synergies.

Here, we addressed these questions by characterizing the temporal evolution of multi-digit kinematics, including the emergence and reorganization of skill-specific synergies, during the early learning of a naturalistic motor skill. We found that complex multi-digit synergies emerge rapidly with practice, are most strongly expressed across early rest periods, and show no evidence of fatigue-related reduction in use. These synergies, composed of small, rapid, and overlapping digit movements, developed gradually over initial training trials and reliably predicted subsequent skill gains, underscoring the evolution of their neural representations in supporting the newly acquired skill.

## Results

We examined kinematic features of early skill acquisition in twenty healthy adults (12 female; mean age 30 years) trained on a well-characterized motor sequence task^14,30–33^. Subjects were tasked with repetitively typing a 5-item keypress sequence displayed on the screen (4-1-3-2-4) as quickly and accurately as possible with 4 digits (index, middle, ring and little fingers) of their left, non-dominant hand. Each trial comprised 10 seconds of practice followed by 10 seconds of rest, yielding a session of approximately 15 minutes length (**Figure 1A**). Motor skill was indexed by the rate of correct sequences per second (cs/s) ^34,35^. The early learning period encompassed the training trials during which 95% of maximum performance was reached ^15,35^, which, at the group level, occurred by trial 12 (**Figure 1B**).

**FIGURE 1:**
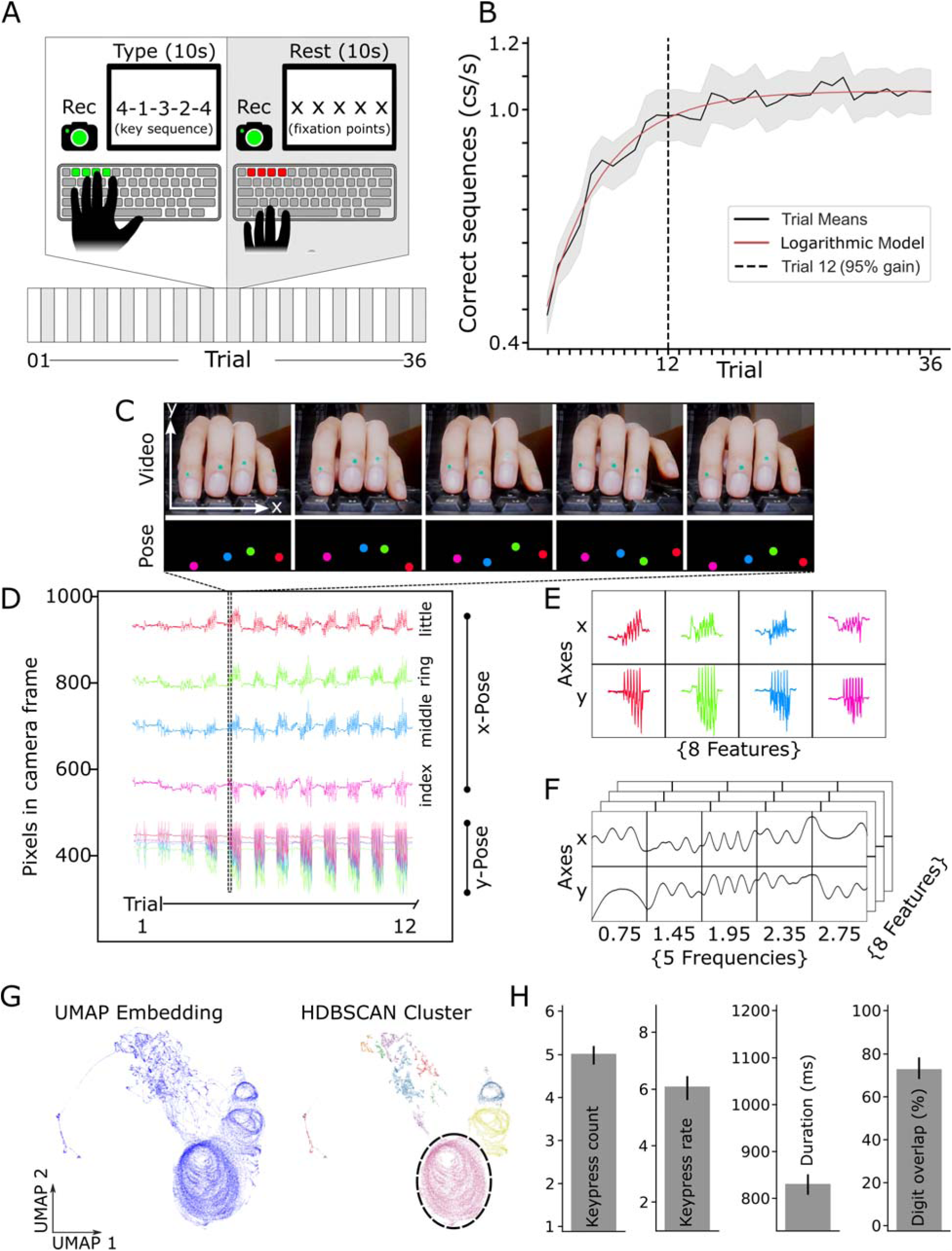
Data analysis. **(A)** Subjects learned a motor sequence over 36 practice trials in a single training session ^32,80^. Each trial alternated between 10 s training (sequence typing) and 10 s rest periods. **(B)** Skill was quantified as the rate of correctly typed sequences per second. Early learning, the training period during which 95% of maximum performance was reached, developed within the initial 12 trials (mean ± 95% CI; logarithmic model fit shown ^15^). The dashed vertical line indicates the trial by which 95% of maximum performance was reached. **(C)** High-definition videos captured from a frontal viewpoint (top) were analysed using markerless pose estimation to track digit positions on each frame (bottom). **(D)** For a representative participant, digit displacement trajectories along the lateral (x) and vertical (y) axes are shown across the first 12 trials, plotted as raw pixel coordinates derived from DeepLabCut. As practice progressed, the amplitude of digit movements increased, becoming especially pronounced by trial 12. **(E)** Digit trajectories during a single practice trial (10sec) in a representative subject yielding 8 features (four digits × two axes). Digit coding as in D. **(F)** We used complex Morlet wavelets cantered on five frequencies known to be relevant during performance of digit movements^39,40^. We decomposed the time series and obtained time-resolved estimates of amplitude and phase. **(G)** Left: Wavelet-transformed digit movements (panel F) were embedded into a two-dimensional space using Uniform Manifold Approximation and Projection (UMAP) shown here for a single participant across the full 36-trial practice session. Each point corresponds to one video frame sampled at 120 Hz. *Right*: Clustering performed on the UMAP embedding using Hierarchical Density-Based Spatial Clustering of Applications with Noise (HDBSCAN) to identify density-defined clusters (behavioural synergies) **(H)** The kinematic properties of a representative synergy cluster (circled in panel G, right) are shown, including the number of keypresses (5 kp), keypress rate (6 kp/s), duration (840 ms), and the proportion of temporal digit overlap within the synergy (75%).

### Characterization of movement synergies

Digit trajectories were recorded at 120 Hz with a front-facing high-speed camera (**Figure 1C, top**). Markerless pose estimation ^36–38^ tracked distal interphalangeal joints, yielding ∼42,500 x–y frames per participant across the full practice session. The x-axis primarily captured in-plane lateral motion (adduction–abduction), whereas the y-axis reflected vertical flexion–extension perpendicular to the keyboard (**Figure 1C-E**). For each practice trial, digit trajectories were represented as multivariate time series comprising eight features (four digits × two spatial axes; **Figure 1E**). Using complex Morlet wavelets centered on five frequencies known to be relevant for digit movements^39,40^, we decomposed the time series to obtain time-resolved estimates of amplitude and phase (**Figure 1F**). For each time point, we formed feature vectors from amplitude and phase across digits and frequencies, embedded them into two dimensions with UMAP (**Methods**) using all practice time points per subject, and applied HDBSCAN (**Methods**) to identify density-defined clusters (behavioural synergies) while labelling low-density points as noise (**Figure 1G**). To quantify training-dependent changes in behavioural synergies, we computed the Jensen–Shannon divergence (JSD) between distributions of cluster occupancy across time, comparing consecutive segments within practice trials (online) and across rest periods (offline) (**Figure S1** shows example cluster distributions and JSD computation). Kinematic features of individual synergies, including keypress count, keypress rate, duration, and inter-digit overlap, were obtained by projecting cluster labels back into the original feature space (**Figure 1H**).

### Emergence of multi-digit synergies during early skill learning

Training was associated with a progressive increase in both the mean keypress count per synergy (**Figure 2A**) and the overall typing rate (**Figure 2B**). In parallel, the duration of individual synergies decreased (**Figure 2C**), while temporal overlap among digit movements increased (**Figure 2D**), together indicating a shift from predominantly isolated single-digit synergies toward more integrated, multi-digit, coarticulated synergies. Indeed, from Trials 1 to 12, one-keypress synergies fell to near extinction, while 2- and 3-keypress synergies rose steeply early on and then declined once skill plateaued. In contrast, 4- keypress synergies increased progressively during early learning (**Figures 2E, S21)**.

**FIGURE 2:**
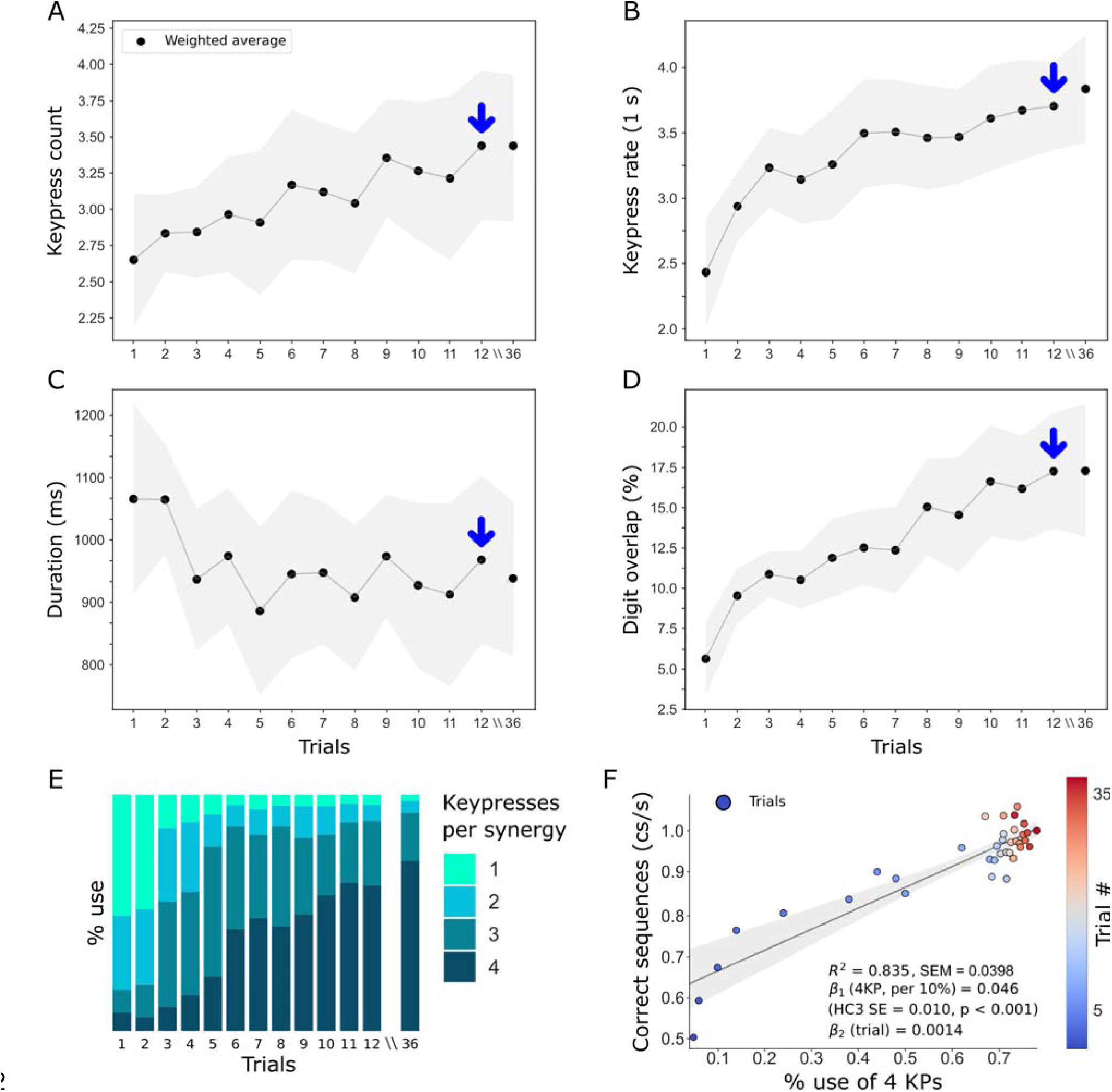
Kinematic properties of motor synergies. **(A)** Mean keypress count (kp/synergy) rose steadily across the early practice trials, attaining an asymptote near trial 12 (blue arrow), after which it plateaued. **(B)** The mean keypress rate (kp/second), normalized by synergy duration, exhibited progressive increases across practice, reflecting steady gains in typing speed. These improvements continued through trial 36. **(C)** The mean duration of individual synergies within each trial decreased rapidly during early practice. **(D)** Digit overlap—defined as the mean proportion of the synergy interval during which multiple digits move concurrently—increased progressively with training. Blue arrows in panels A–D indicate trial 12, the point at which participants had achieved approximately 95% of their maximum performance, marking the end of early learning ^15^. **(E)** Stacked bar plots show the proportional deployment of synergies comprising one, two, three, or four keypresses across training. Note that with practice, participants gradually reweighted their repertoire from early reliance on single/double keypress synergies (light blues) to preferential use of synergies containing three and four keypresses (darker blues). **(F)** Linear regression was used to quantify the relationship between the proportion of four-keypress synergies and skill (cs/s), with each point representing a single trial and color-coded by trial number (1–36). Across practice, increased reliance on four-keypress synergies closely paralleled improvements in skill (R² = 0.83, p < 0.001; SEM = 0.0398).

Synergy structure changed predominantly during Trials 1–12 (**Table 1**; Stouffer’s p ≈ 6 × 10⁻¹□). In the later phase (Trials 12–36), the corresponding effect did not reach significance at the group level (p = 0.107; **Table 2**). This non-significant result was not accompanied by evidence for stability, as individual JSD values remained non trivial and an equivalence test (δ = 0.15 JSD) did not support equivalence. Critically, the % use of 4-keypress synergies correlated with the magnitude of skill learning (R² = 0.83, p < 0.001; SEM = 0.0398; **Figures 2F and Figures S2 D, E)**. Collectively, these findings indicate that early practice rapidly prunes the motor repertoire toward temporally compressed, overlapping, high-order multi-digit synergies predictive of the magnitude of the acquired skill.

**Table 1:**
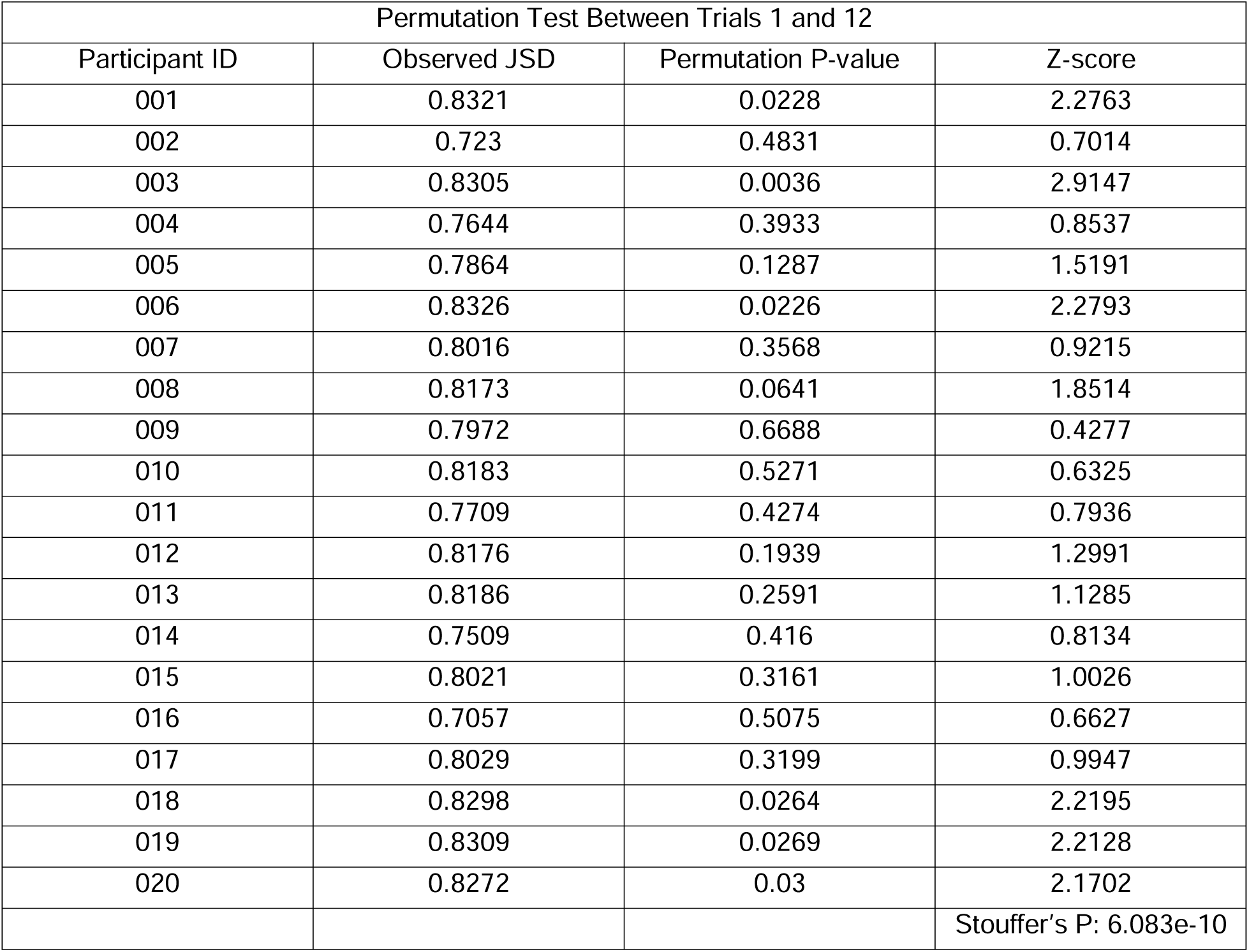
Permutation tests comparing synergy distributions between early and late training trials. Jensen–Shannon Divergence (JSD) quantified differences in synergy distributions between pairs of trials for each participant. Permutation tests assessed the statistical significance of observed JSD values (permutation p-values), which were converted into Z-scores. Across participants, Z-scores were combined to produce a group-level significance using Stouffer’s method (Stouffer’s p). Table 1 (top) compares Trials 1 and 12, revealing a significant change in synergy distributions across participants (Stouffer’s p = 6.083e-10). Table 2 (middle) compares Trials 12 and 36, indicating no significant change across participants (Stouffer’s p = 0.1065).

**Table 2:** Permutation tests comparing synergy distributions between early and late training trials. Jensen–Shannon Divergence (JSD) quantified differences in synergy distributions between pairs of trials for each participant. Permutation tests assessed the statistical significance of observed JSD values (permutation p-values), which were converted into Z-scores. Across participants, Z-scores were combined to produce a group-level significance using Stouffer’s method (Stouffer’s p). Table 1 (top) compares Trials 1 and 12, revealing a significant change in synergy distributions across participants (Stouffer’s p = 6.083e-10). Table 2 (middle) compares Trials 12 and 36, indicating no significant change across participants (Stouffer’s p = 0.1065).

| Permutation Test Between Trials 12 and 36 |  |  |  |
| --- | --- | --- | --- |
| Participant ID | Observed JSD | Permutation P-value | Z-score |
| 001 | 0.8055 | 0.3724 | 0.8921 |
| 002 | 0.6119 | 0.8466 | 0.1935 |
| 003 | 0.4169 | 0.985 | 0.0188 |
| 004 | 0.4043 | 0.9645 | 0.0445 |
| 005 | 0.4633 | 0.9721 | 0.035 |
| 006 | 0.5681 | 0.7679 | 0.2951 |
| 007 | 0.4554 | 0.931 | 0.0866 |
| 008 | 0.4066 | 0.989 | 0.0138 |
| 009 | 0.8326 | 0.0112 | 2.5355 |
| 010 | 0.6235 | 0.8024 | 0.2502 |
| 011 | 0.3905 | 0.9668 | 0.0416 |
| 012 | 0.6548 | 0.7542 | 0.3131 |
| 013 | 0.7429 | 0.3663 | 0.9034 |
| 014 | 0.6653 | 0.6777 | 0.4157 |
| 015 | 0.7945 | 0.4672 | 0.727 |
| 016 | 0.3753 | 0.9754 | 0.0308 |
| 017 | 0.4596 | 0.8104 | 0.2399 |
| 018 | 0.3812 | 0.9785 | 0.027 |
| 019 | 0.4742 | 0.9174 | 0.1037 |
| 020 | 0.4269 | 0.9591 | 0.0512 |
|  |  |  | Stouffer's P: 1.065e-1 |

### No detectable fatigue-related reduction in expert synergy use

We next asked whether accumulating fatigue with continued practice^41^ might influence this kinematic transformation. Fatigue-related motor slowing^42^ could result in the progressive attrition of multi-digit synergies, either within individual 10-s practice trials or cumulatively across the 36-trial session. Contrary to these predictions, once multi-digit synergies were established by the end of early learning, their expression showed no decline within individual trials at the end of training (when fatigue could be expected, **Figure S3**) nor across the entire training session **(Figure 2E)**. Thus, early learning of a naturalistic skill proceeds without detectable fatigue-related reduction in use of multi-digit synergies.

### Higher-Order Multi-Digit Synergies Emerge Across Rest Intervals Interleaved Practice

Prior work indicates that early gains in challenging, naturalistic motor skills may accrue across brief rest intervals (“micro-offline”) to a larger extent than during active practice (“micro-online”)^15^. Here, we replicated this observation. On average, micro-online changes were negligible (relative to 0, no change, mean = −0.297 ± 0.325 cs/s; two-tailed one-sample t-test, t₁₉ = −0.53, p = 0.60), whereas micro-offline gains were robust (mean = 1.006 ± 0.317 cs/s; two-tailed one-sample t-test, t₁₉ = 4.61, p = 1.92 × 10⁻□). Micro-offline gains significantly exceeded micro-online changes (paired t-test, t₁₉ = 3.68, p = 0.0016) and were statistically indistinguishable from total early learning (mean = 0.709 ± 0.055 cs/s; paired t-test, t₁₉ = 1.86, p = 0.079).

Changes in HDBSCAN-derived synergy maps (**Figure 1G**) developed predominantly across rest (mean = 0.596; SEM ±0.026) rather than practice (mean = .316; SEM ±0.028) intervals (**Figure 3A**). JSD divergence across rest intervals predicted early learning (R² = 0.711, p < 0.001; **Figure 3C**), whereas JSD divergence during practice bouts did not (R² = 0.104, p = 0.166; **Figure 3B**). This result was robust across different hyperparameter settings, yielding comparable effect sizes and significance levels (**Figure S3 B, C, D**).

**FIGURE 3.**
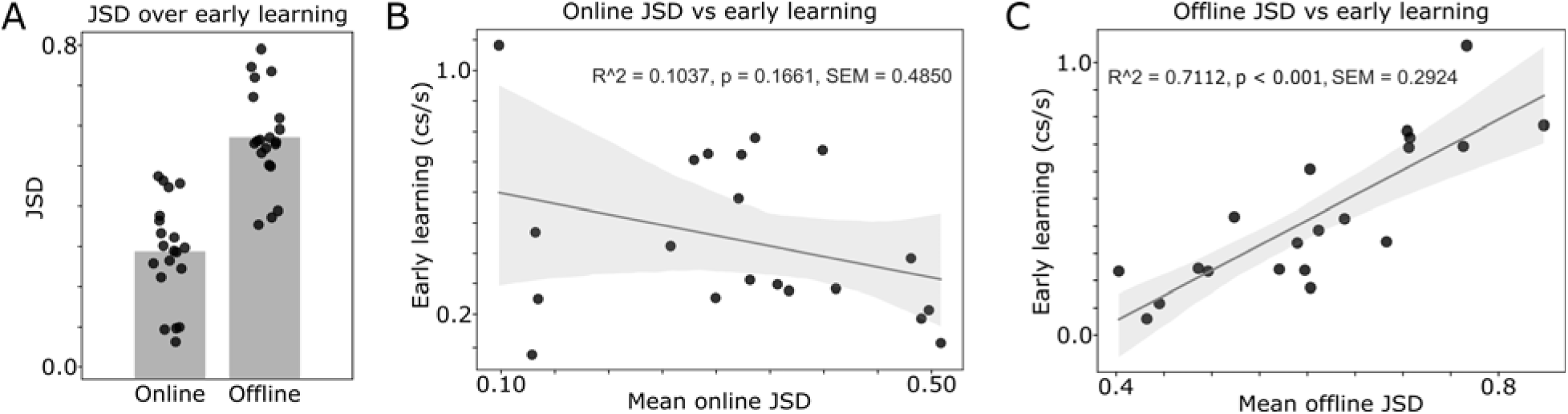
HDBSCAN-derived synergy maps reorganize more during rest than during practice in early learning. **(A)** During early learning (trials 1–12), Jensen–Shannon divergence (JSD) between consecutive synergy maps was greater across rest intervals than across practice intervals, indicating stronger reorganization during brief pauses. **(B)** Changes in synergy maps across practice intervals (online JSD) did not reliably predict early learning (R² = 0.104, p = 0.166, SEM = 0.485). **(C)** By contrast, changes in synergy maps across rest intervals (offline JSD) strongly correlated with early learning (R² = 0.711, p < 0.001, SEM = 0.292), suggesting that key aspects of synergy reconfiguration supporting early performance gains occur predominantly offline.

Together with the absence of a detectable fatigue effect on multi-digit synergy use during acquisition of this challenging naturalistic skill, these findings are consistent with the view that the transition to higher-order multi-digit coordination may be shaped predominantly by micro-offline processes unfolding across brief rest intervals.

### Synergies subserving early learning of a naturalistic skill

Next, we quantified the emergence, decay, and overall prevalence of 1–5-keypress synergies at two timescales, within trials and across the full course of training, using broad prevalence bins for visualization (**Figure 4**, representative single subject; **Figure 5**, group data). Because prevalence varied continuously and showed no clear natural boundaries, synergies were grouped by session-wide percent usage into four heuristic categories: <5%, 6–20%, 21–49%, and >50%. These bins provided sufficient observations within each category while distinguishing rarely expressed from more prevalent synergies.

**Figure 4:**
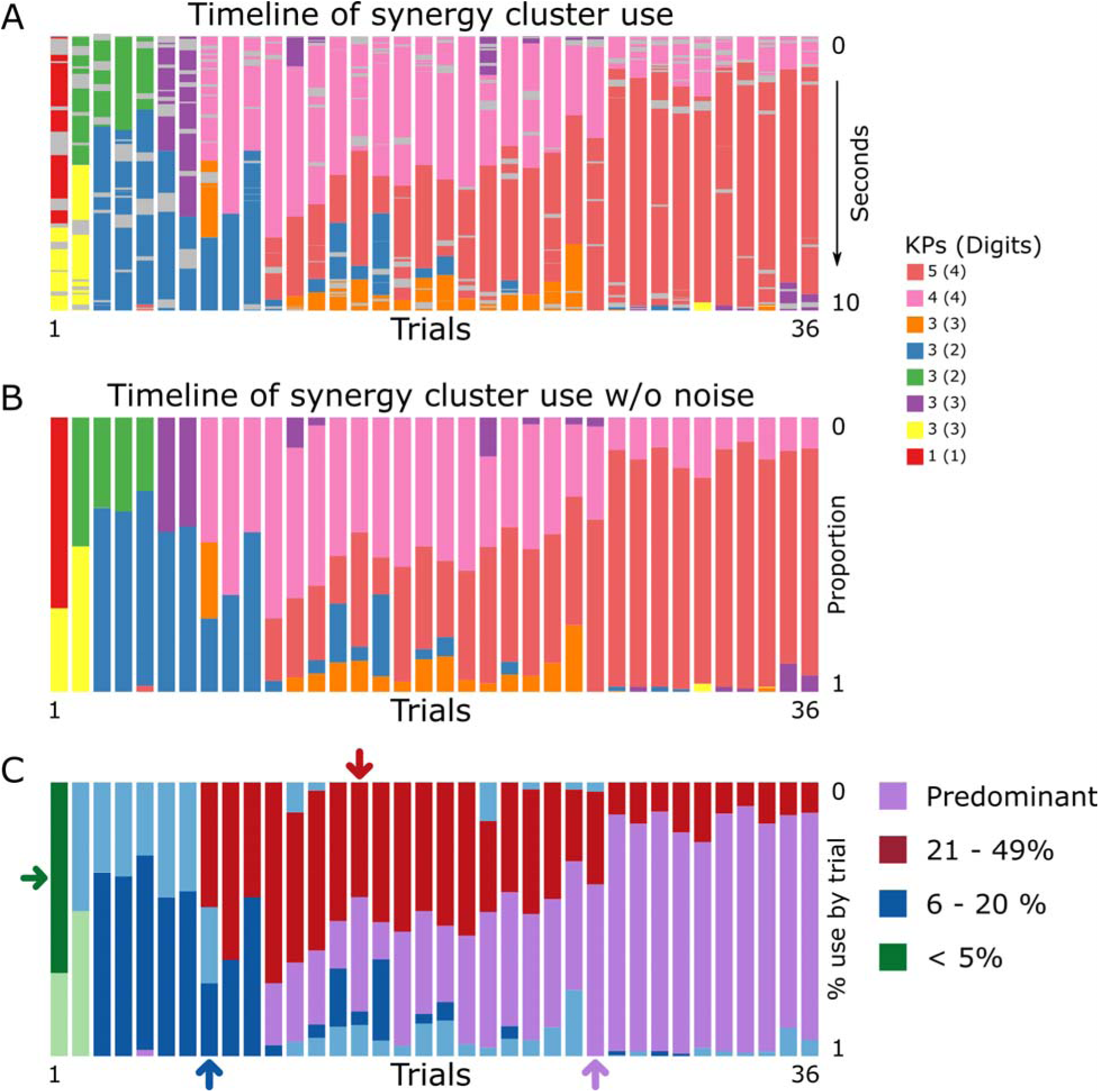
Emergence of expert synergies during early learning in a representative subject. **(A)** Bar plot depicting the temporal distribution of synergy clusters across each 10-s practice interval over 36 trials in the same representative participant depicted in **Fig 1G**. Gray segments mark epochs within a trial in which no stable synergies were identified (noise). Across training, a single predominant synergy (purple) emerges and progressively dominates behavior, illustrating the stabilization of a preferred coordination pattern with practice. **(B):** Timeline of synergy cluster occurrences (excluding noise), normalized to trial duration (0–1) for the same representative participant. This bar plot parallels panel A but omits gray noise epochs, showing only the sequence of identified synergies across each 10-s practice interval (n = 36 trials). This noise-free, time-normalized representation was used for all subsequent JSD-based statistical analyses (**Figure S1**). The number of keypresses and digits per synergy is indicated on the right. Early in training, synergies predominantly comprise single or double digits, whereas later-emerging, dominant synergies recruit multiple digits, reflecting the development of higher-order coordination. **(C)** Synergies were subsequently classified according to their overall percentage of use across all training trials (>50%, 21–49%, 6–20%, and <5%). Several distinct temporal profiles emerged in this subject. Some synergies were expressed almost exclusively in the very first trial(s) (e.g., green arrow, “novice”), whereas others appeared transiently during mid-practice (e.g., dark blue arrow, “exploratory”). Certain synergies were expressed predominantly at trial onset as performance approached asymptote (e.g., red arrow, “trial initiation”), while others emerged around mid-training and became increasingly dominant toward the end of practice (e.g., purple arrow, “expert”). This qualitative pattern, illustrated here for a single participant, was also observed at the group level (Fig. 5).

**Figure 5.**
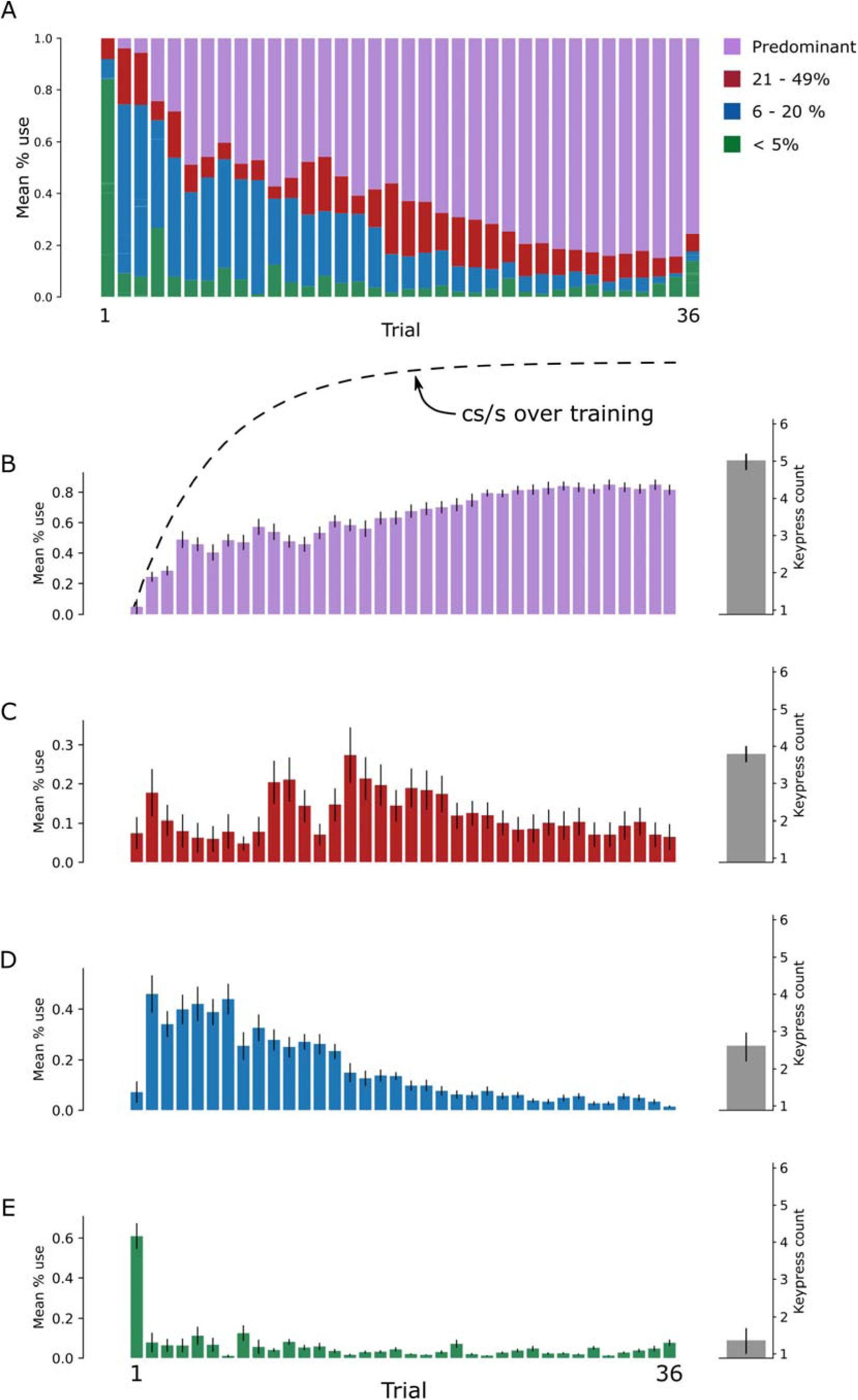
Synergy types emerging during early learning of a challenging naturalistic skill. **(A)** Synergies were classified according to their overall percentage of use across all training trials (>50%, 21–49%, 6–20%, <5%). Group-level stacked bars show the average percent use of each synergy class across trials (1–36). **(B–E)** Group-averaged percent use (± SEM) for each synergy type across trials and subjects. The learning curve is overlaid in panel **B** (dashed line) for reference. Gray bars to the right of each panel indicate the mean number of keypresses encompassed by each synergy type (± SEM). At the group level, the pattern observed in the single-subject example (Fig. 4) is preserved: predominant 5-keypress synergies late in training (**B**, “expert”), trial initiation 2-4 keypress synergies expressed relatively uniformly across training (**C**, “trial initiation”), synergies engaging variable number of keypresses that emerge and fade mid-training (**D**, “exploratory”), and early 1–2-keypress synergies largely confined to the first trial (**E**, “novice”).

Synergies used for <5% of training time appeared almost exclusively in the earliest trial(s), comprised 1-2 keypresses/synergy and then disappeared (green arrow in **Figure 4C**; **Figure 5E**). Synergies expressed for 6–20% of training time, emerged and faded mid-practice and likewise comprising 1–2 keypresses per synergy (dark blue arrow in **Figure 4C**; **5D**). Synergies expressed for 21–49% of training time included a type that occurred reliably at the start of trials as performance approached asymptote, persisted through session end, and encompassed 3-4 keypresses per synergy (red arrow in **Figure 4C; 5C**). Finally, synergies present for >50% of training time comprised the “predominant/expert” patterns, which arose mid-training, became dominant toward the end of practice, and could encompass up to 5 keypresses per synergy (purple arrow in **Figure 4C**; **5B**).

The most prevalent composition of the predominant expert multi-digit synergies observed in the last practice trial was “4–1–3–2” in 52% of subjects (**Figure S4**, **S5 and S6**). The second most prevalent was “4–1–3–2-4” in 21% of *subjects*, the full trained sequence, which incorporate a repeated “4” spanning the sequence boundary (last keypress of one sequence followed by the first keypress of the next) to gain speed^43^. Next was “4-4-1” present in 16% of the subjects, also taking advantage of the 4–4 double tap. Finally, “4-4-1-3-2” was the predominant synergy in 11% of the subjects. Note that the last two expert synergy types crossed individual sequence boundaries. Together, these findings indicate that skilled performance in this task is supported by a relatively small repertoire of expert multi-digit synergies some of which exploit boundary double taps and possibly sequence fractionation (chunking) ^44^.

## Discussion

Here we show that complex, multi-digit synergies emerge rapidly after practice onset and increase progressively during the early phases of naturalistic skill learning, with their most prominent development occurring across intervening rest breaks. The synergies that supported acquisition of the trained skill were characterized by small, rapid, and overlapping digit movements that gradually crystallized over early trials and robustly predicted subsequent performance gains. Notably, we did not observe fatigue-dependent reductions in multi-digit synergy use.

### High-Order Multidigit Synergies Underlie Improvements in Naturalistic Skill

Motor learning, from everyday skills such as typing to highly trained behaviors like musical performance, relies on the production of temporally precise action sequences that form the basis of fine motor expertise ^16,44–46^. These sequences are not controlled muscle by muscle, but through coordinated motor synergies, making their reorganization central to skilled performance. Accordingly, examining how motor synergies reorganize with learning provides a direct window into the neuromotor processes that support the acquisition, refinement, and generalization of skilled actions.

To address this question, we recorded movements with a high-speed camera while subjects learned a new sequential motor skill. Markerless pose estimation software ^47^ and a dimensionality reduction strategy^47^ allowed us to characterize synergies (unique combinations of covarying four-digit movements over one to five keypresses), as the new skill was learnt. Multi-digit synergies are the ultimate expression of efficient modular control strategies used by the central nervous system to execute sequential skills, like playing musical instruments^48–53^. We observed that synergy morphology evolved remarkably quickly, changing within seconds of training onset. Early practice rapidly sculpted the motor repertoire toward temporally compressed, overlapping, higher-order multi-digit synergies, whose dominance from approximately the twelfth practice trial onward, when performance near plateaued, robustly predicted the degree of skill acquisition and likely reflected an expansion of underlying memory capacity^28^.

Synergies involving single digits, which predominated during the earliest trials, progressively transitioned into coordinated multi-digit patterns as training advanced. This rapid reorganization of synergy structure closely mirrored the time course of behavioral improvement. This transition may parallel the rapid shift from stimulus-bound responding to internally generated sequence production, in which actions are guided by an internal memory representation rather than discrete external cues, as is typical of reaction-time–based paradigms^54^. The swift emergence of this internally driven mode of control suggests that developing motor kinematics, associated with an expanding skill memory ^28^, may become primary constraints on behavior.

### Mechanisms Underlying the Formation of Expert Synergies

How might multi-digit synergies emerge preferentially across rest rather than during practice? We found that behavioral improvements occurred primarily across the brief rest periods interleaved with practice (i.e., micro-offline gains), consistent with prior reports ^15,19,27,34,55–57^. In the context of challenging skills, micro-offline skill gains are up to four times larger than those identified following overnight sleep^15^, are reproducible ^57–61^ and persist even when practice bouts are shortened to as little as 5 seconds, arguing against recovery from performance fatigue as their primary source ^58^. It has been proposed that this form of micro-offline gains represent a form of memory consolidation^15,54,62^, a view supported by mechanistic work^34^ ^60,61^ ^62^.

Our results extend prior work by demonstrating that motor synergy maps can reorganize across rest periods, revealing a previously unrecognized neural–motor correlate of micro–offline learning. More broadly, our findings suggest that practice and rest make distinct yet complementary contributions to naturalistic skill learning. Practice itself may primarily encode elemental action representations and provide the feedback necessary to optimize movement kinematics^63,64^. Rest intervals interleaved with practice may promote the binding of individual action components into higher-order synergies, effectively compressing the action space and freeing capacity for further online learning^28^. This interpretation accords with prior accounts suggesting that the relative contributions of online and offline processes to learning are shaped by task configuration^28^ and context^65^, and that these processes are complementary rather than competitive^18^.

Although our design followed published guidelines to minimize it ^42^, accumulating fatigue with continued practice ^41^ could, in principle, reduce the use of expert multi-digit synergies either within individual 10-s practice trials or cumulatively across the 36-trial session ^66,67^. We found no evidence this occurred. Once multi-digit synergies were established by the end of early learning, their use showed no decline within trials at the end of training (when fatigue would be maximal in the session, **Figure S3**) or across the entire training session **(Figure 2E)**. Thus, across these two independent measures we found no evidence that use of expert synergies during early learning of a challenging naturalistic skill is reduced by motor or cognitive fatigue.

### Do Distinct Synergy Subtypes Subserve Early Naturalistic Learning?

To aid interpretation, we classified synergies by the proportion of training time during which they were expressed. These categories were intended as descriptive rather than as discrete synergy classes. Because prevalence varied continuously across the session and showed no clear natural boundaries, we grouped synergies into four heuristic bins (<5%, 6–20%, 21–49%, and >50%) that balanced interpretability with adequate sampling. This framework distinguished relatively rare from more predominant synergies and allowed us to consider whether expression frequency may relate to distinct functional roles during skill acquisition. A set of synergies expressed for <5% of total training time (1–2 keypresses/synergy) appeared predominantly in the first few trials and then vanished (**Figure 4C**, green arrow; **Figure 5E**). These “novice” synergies likely reflect provisional kinematic solutions to the demands of a newly acquired skill, which are rapidly supplanted by higher-order, multi-digit synergies as learning unfolds. A second subgroup consisted of synergies expressed for 6–20% of training time (1–2 keypresses per synergy), which emerged and faded mid-practice (**Figure 4C**, dark blue arrow; **Figure 5D**). These “exploratory” synergies may reflect kinematic experimentation during the transition from stimulus-driven to internally generated execution of the naturalistic skill. Synergies expressed 21–49% of training time (1–4 keypresses per synergy) appeared consistently at trial onset as performance neared asymptote and remained present until the end of the session (**Fig. 4C**, red arrow**; 5C**). These “trial-initiation” synergies may index the contribution of advance planning to the earliest phase of trial performance, a possibility consistent with estimates of the preplanning capacity of 2-4 keypresses in sequence learning^68–70,71^.

Synergies expressed for >50% of total training time (up to five keypresses per synergy) constituted the “predominant/expert” class: they first appeared around mid-training and progressively came to dominate behavior toward the end of practice (**Figure 4C**, purple arrow; **Figure 5B**). The dominant features of these expert synergies were optimized speed–accuracy for repeated taps on the same key (or double taps)^43^ and systematic fractionation of the trained sequence into preferred chunks^72–74^. This reorganization of skill synergies likely supports the early integration of fragmented, independent movements into highly coordinated, multi-digit actions, effectively reducing the functional degrees of freedom in expert performers ^5,6,75^.

The tight coupling between a small set of expert multi-digit synergies and high skill levels raises an intriguing translational question: could explicit training of these expert synergies complement conventional practice and accelerate learning in domains such as sport, musical performance, or post-stroke neurorehabilitation^76^? More broadly, a deeper understanding of the kinematic mechanisms that underlie skill acquisition may inform the design of more effective assistive and rehabilitative technologies, for example by guiding synergy-based control strategies in neuroprosthetics or robotics ^7,77^.

### Limitations and Future Directions

Several limitations of the present work suggest fruitful avenues for future research. First, employing multi-camera motion capture would permit full three-dimensional kinematic reconstruction, enabling a more comprehensive characterization of movement structure than was possible here. Second, extending synergy analyses beyond distal digit segments to include additional digits and proximal effectors that support manual function, such as the wrist, arm, and shoulder, may uncover further kinematic contributions to skilled performance and clarify how distal and proximal control are integrated. Finally, future studies should test whether the kinematic mechanisms underlying sequence learning generalize to multi-limb coordination, postural control, and other forms of motor learning, an issue central to evaluating their broader relevance for neurorehabilitation of skilled function.

## Authors Contributions

L.G.C. and E.R.B. conceived of the presented idea. W.K., E.R.B. and L.G.C. designed the experiment. W.K., R.F., G.R.R. and M.H. carried out the experiment. W.K. developed and tested the analysis pipeline. W.K. L.G.C. and S.B. carried out the analyses. W.K., S.B. and L.G.C. wrote the manuscript. W.K., S.B. and L.G.C. reviewed and edited the manuscript.

## Resource Availability

### Lead Contact

William D. Kistler.

### Materials availability

This study did not generate new unique reagents.

### Data and Code Availability

Data used in this study (subject to participant consent) will be made available upon reasonable request to the lead contact. All custom code used for processing, embedding, clustering, and statistical analysis is publicly available and archived at Zenodo [https://zenodo.org/records/10717889].

## Acknowledgments

This research was supported by the Intramural Research Program of the National Institutes of Health (NIH). The contributions of the NIH author(s) are considered Works of the United States Government. The findings and conclusions presented in this paper are those of the author(s) and do not necessarily reflect the views of the NIH or the U.S. Department of Health and Human Services. This research was also supported by the NIMH-UCL Graduate partnership program.

## STAR Methods

### Key Resources Table

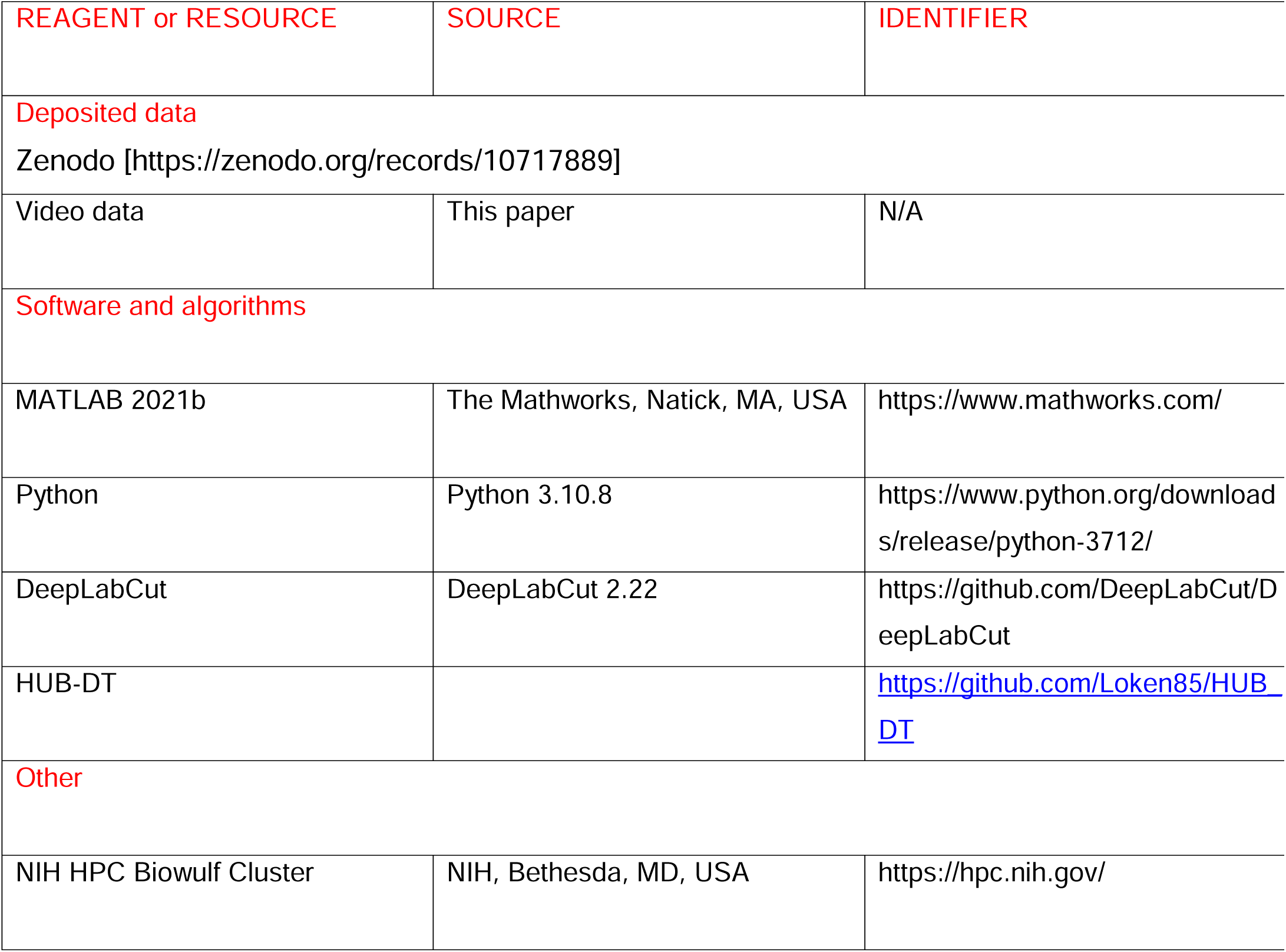

### EXPERIMENTAL MODEL AND SUBJECT DETAILS

#### PARTICIPANTS

Twenty right-handed, neurologically healthy adults (12 women; mean age = 29.2 ± 2.12 years, SD) completed participation under a protocol approved by the NIH Combined Neuroscience Institutional Review Board. Although active musicians were excluded to minimize the influence of extensive fine motor training, we did not formally quantify typing experience or keyboard proficiency. Seventeen additional participants were excluded due to technical issues with video capture, yielding a final dataset of 20 participants. The sample size was determined *a priori* based on our primary planned analysis, a linear regression testing whether keypress sequence skill learning is predicted by the prevalence of 4-digit synergies. Because no published data were available to estimate the expected strength of this specific relationship, we used the average R^2^-value from two previously reported associations between learning in this task and neural activity features (neural replay density^34^ and representation distance^54^) as a proxy. We assumed that the association between two behavioral measures would be at least as large as this proxy estimate. For α = 0.05, 99% power (1 - β = 0.99) and an anticipated large effect-size (R^2^ = 0.5097), a sample-size of 20 participants was determined to be sufficient for detecting a significant linear relationship between skill learning and 4-digit synergy prevalence.

#### TASK

Participants performed a 5-digit sequence-typing task (41324) with their non-dominant (left) hand on a standard 104-key QWERTY keyboard. The little, ring, middle, and index fingers were mapped to numeric keys 1 through 4, respectively; the thumb was not used. Each trial consisted of 10 s of practice followed by 10 s of rest, and participants completed 36 such trials within a single session. Throughout practice, the target sequence was continuously displayed, and each keypress elicited an asterisk, providing real-time feedback on sequence progression but not on accuracy. During rest periods, the sequence was replaced by five “X” characters to discourage overt or covert rehearsal. The task was administered, and all responses were recorded, using PsyToolkit (www.psytoolkit.org).

### DATA ANALYSIS

#### POSE ESTIMATION

Digit kinematics were captured at 120 frames/s using a high-speed camera positioned in front of the left hand (**Figure 1C**, top). Markerless pose estimation ^36,37^ was used to localize the distal interphalangeal joints of the four active fingers (**Figure 1C**, top) ^38^, yielding approximately 43,000 frames of x–y pose data per participant over the full practice period (**STAR Methods**). The x coordinate primarily indexed lateral motion within the keyboard plane (adduction–abduction), whereas the y coordinate reflected vertical displacement perpendicular to the keyboard surface (flexion–extension; **Figure 1C–E, STAR Methods**).

Pose estimation was implemented in DeepLabCut v2.2.2 (Python 3.10.8) with GPU acceleration via CUDA Toolkit 11.2 and cuDNN 8.2. Sixty frames per participant (1,200 total) were manually annotated for the x,y positions of all four fingers. A ResNet-50 backbone was trained for 400,000 iterations using DLC default settings (three color channels, pairwise terms enabled, unsupervised refinement disabled). For a single shuffle, training and test errors were 2.12 and 2.18 pixels, respectively (image resolution: 1280 × 720) (**Figure S6**).

Framewise predictions with likelihood < 0.95 were flagged as low confidence and subjected to visual inspection. Missing samples were linearly interpolated when gaps were ≤ 3 consecutive frames; longer gaps were excluded from further analysis. No temporal smoothing or filtering was applied to the resulting trajectories prior to downstream analyses.

#### SKILL MEASUREMENT

Motor skill was operationalized as the number of correct sequences produced per second (cs/s). A sequence was scored as correct if it matched the canonical pattern (41324) or any of its circular permutations (e.g., 13241, 32414). For each 10 s trial, we computed the average instantaneous cs/s. To characterize learning over time, cs/s data were fit with an exponential growth function:

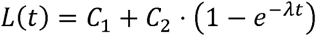

where (C_1_) denotes baseline performance, (C_2_) the asymptotic learning gain, and (\lambda) the rate constant. Parameters were estimated using constrained nonlinear least-squares (trust-region-reflective algorithm). Cumulative learning was quantified by integrating (L(t)) over practice time and normalizing trial-wise gains by the total area under the curve. Across participants, 95% of total gains (period defining early learning) were achieved by Trial 12.

#### ONLINE AND OFFLINE LEARNING CONTRIBUTIONS

Microscale changes in skill performance were decomposed into online and offline components using correct sequences per second (cs/s) as the behavioral metric. Micro-online gains were defined as the within-trial change in cs/s, computed as the difference between the average performance in the first and last second of each 10-s practice trial. Micro-offline gains were defined as the between-trial change in cs/s, computed as the difference between the last second of a given trial and the first second of the subsequent trial. These measures were calculated over the first 12 trials, corresponding to the early learning phase. Cumulative micro-online and micro-offline changes were then summed across this interval to estimate their relative contributions to overall skill acquisition.

To characterize structural changes in motor behavior, we examined trial-level synergy distributions based on HDBSCAN-derived cluster labels assigned to each video frame. For each 10-s trial, we extracted the first and last 1-s windows (120 frames each at 120 fps) and quantified the composition of behavioral clusters (synergies) within each window as a normalized histogram, excluding frames labeled as noise. Jensen–Shannon divergence (JSD) was computed between the first and last second of each trial to index online synergy reorganization, and between the last second of one trial and the first second of the next to index offline reorganization. This approach enabled a direct comparison between kinematic reconfiguration and skill change at the microscale.

#### POSE TRANSFORMATION AND SYNERGY EXTRACTION

To incorporate temporal structure into the pose data, we applied Morlet wavelet transforms to the x- and y-coordinates of each digit at five frequencies (0.75, 1.45, 1.95, 2.35, and 2.75 Hz), chosen to span the range of rhythmic, voluntary finger movements observed in pilot recordings. This yielded a time–frequency representation for each frame in which each digit’s position was decomposed into oscillatory amplitudes at each frequency. The aim was to capture behavioral variation across multiple timescales and to allow for coordination between digits operating at potentially different frequencies—for example, distinct rhythmic patterns expressed by the pinky and index fingers within the same behavioral segment. The resulting feature vector for each frame comprised 40 dimensions (4 digits × 2 coordinates × 5 wavelets).

We performed dimensionality reduction using Uniform Manifold Approximation and Projection (UMAP), as implemented in HUB-DT ^78^. For each participant, frame-wise feature vectors comprising wavelet-derived amplitude and phase across four digits and frequencies were embedded into a two-dimensional space (n_neighbors = 40, n_components = 2, min_dist = 0.1, metric = Manhattan). All time points from the practice session were included in the embedding. The resulting two-dimensional representation was used as input to HDBSCAN for density-based clustering of behavioural synergies. ^78^

#### CLUSTERING OF BEHAVIOURAL DATA

To identify motor synergies, we applied HDBSCAN (Hierarchical Density-Based Spatial Clustering of Applications with Noise) to the two-dimensional UMAP embedding of the wavelet-transformed pose data ^78^. Each point in this embedding corresponded to a single video frame, representing the instantaneous kinematic state derived from the wavelet-transformed features. HDBSCAN grouped points into clusters based on local density (minimum cluster size = 200, minimum samples = 30, cluster selection = “leaf”, cluster_selection_epsilon = 0.14), labelling sparsely populated regions as noise (label −1). Each resulting cluster was treated as a discrete behavioral unit, or motor synergy, whereas noise labels captured transitional or non-repetitive movements.

The resulting labels provided frame-level, fully unsupervised classifications of movement, enabling objective segmentation of behavioral structure across training. To characterize each synergy, we computed mean wavelet amplitude profiles across all frames within a cluster. These profiles summarized the dominant spatial and frequency features of each motor pattern, including digit-specific rhythmic contributions. For each trial, we computed the proportion of frames assigned to each synergy as a normalized histogram (summing to 1). Frames labelled as noise were excluded from this quantification. These normalized synergy distributions served as the kinematic fingerprint of each trial and were used in all between-trial comparisons.

### JENSEN-SHANNON DIVERGENCE AND PERMUTATION TESTING

To quantify how motor synergy distributions changed with learning, we computed the Jensen–Shannon divergence (JSD) between behavioral label distributions at different trial time points. JSD was chosen because it is symmetric, bounded, and robust to zero probabilities ^79^. Further, JSD captures proportional changes across all labels in a scale-invariant and interpretable way, avoids directional bias, and uses smoothing that prevents divergence due to missing cluster labels.

For each participant, synergy labels within a trial were converted into a probability distribution by computing the relative frequency of each cluster. For each pair of trials, JSD was then calculated between these distributions to index changes in synergy composition. To assess whether observed JSD values exceeded chance, we used a nonparametric permutation test: cluster labels were randomly reassigned 100,000 times between the two distributions, recomputing JSD on each iteration to generate a null distribution. The empirical p-value was defined as the proportion of permuted JSD values greater than or equal to the observed value, preserving marginal structure while randomizing label correspondence.

For group-level inference, individual p-values were transformed to z-scores using the inverse standard normal distribution and combined across participants using Stouffer’s method, yielding a single aggregate z-statistic and Stouffer’s p-value.

### QUANTIFICATION AND STATISTICAL ANALYSIS

All modeling and statistical analyses were conducted in Python 3.10.8 using NumPy, SciPy, UMAP-learn, HDBSCAN, and scikit-learn. Exponential fits were obtained with scipy.optimize.curve_fit using boundary-constrained nonlinear least squares. JSD was computed using log-sum averaging, and permutation tests were implemented with custom Python code. Individual z-scores were combined at the group level using Stouffer’s method. Statistical significance was defined as α = 0.05. All code and configuration files are available on Zenodo [https://zenodo.org/records/10717889].

## Supplementary figures

**Figure S1.**
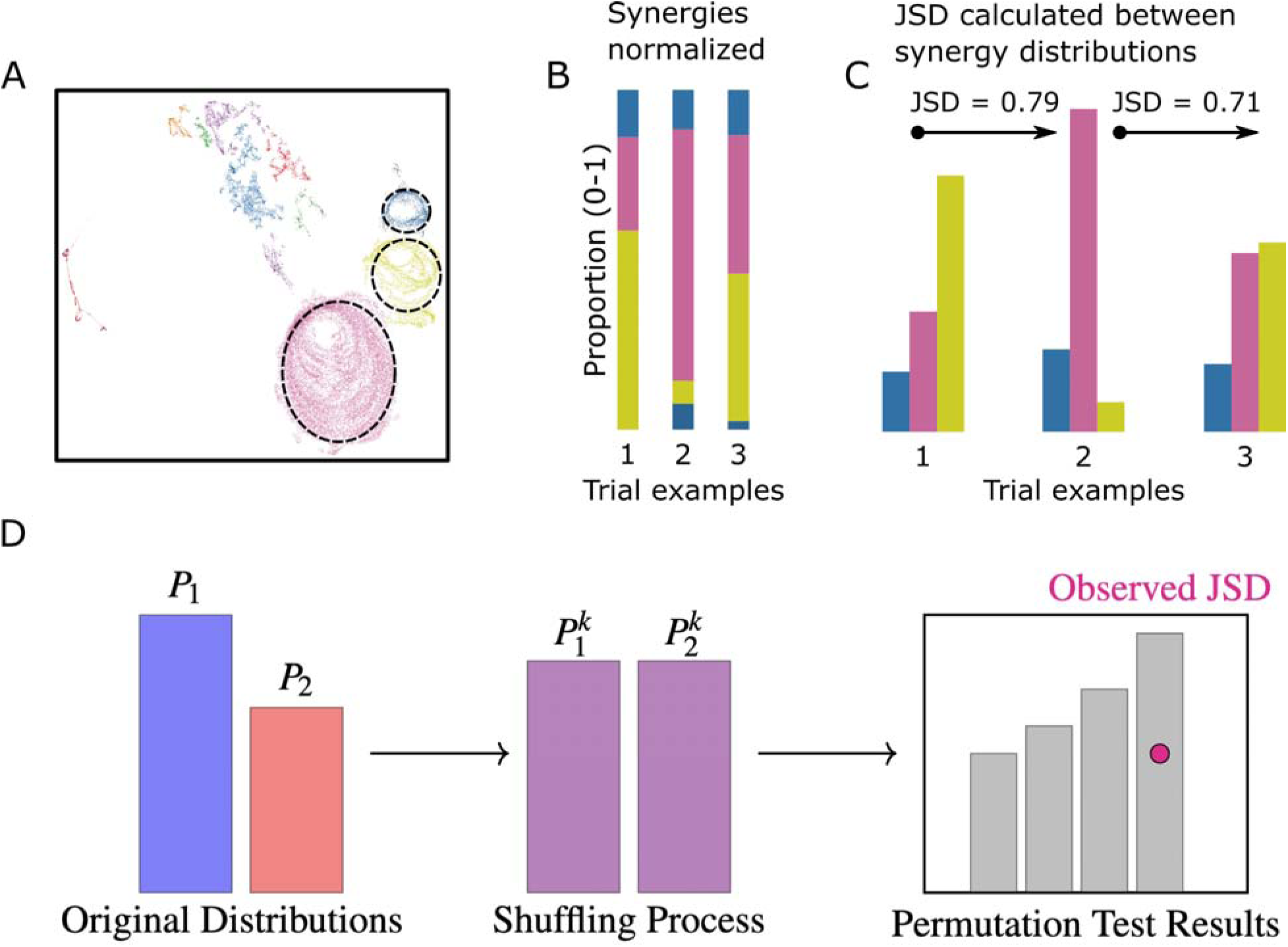
related to STAR Methods. **(A)** UMAP embedding of an illustrative participant’s behavioral data (same as in Figure 1G), highlighting selected multi-digit synergy clusters. **(B)** Normalized synergy distributions for three illustrative trials (noise removed). Each vertical bar depicts the proportional contribution of each synergy within a trial. **(C) Top**. Synergy data from (B) reorganized to enable direct comparison of synergy proportions across trials. Bars of the same color correspond to the same synergy label, and bar height indicates that synergy’s proportion within a trial. When a synergy occurred multiple times in a trial, its contributions were summed (e.g., blue synergy in trials 2 and 3). Jensen–Shannon divergence (JSD) values quantify differences between synergy distributions for each trial pair (e.g., trials 1 vs. 2, 2 vs. 3), providing a numerical index of behavioral reorganization during learning. **Bottom**. Conceptual overview of the permutation test used to assess the statistical significance of observed JSD values. Original synergy distributions (left) are randomly reassigned across synergy labels (center) to generate a null distribution of JSD values (right). The empirical JSD (pink dot) is then evaluated against this null distribution to determine whether the observed behavioral change exceeds chance.

**Figure S2.**
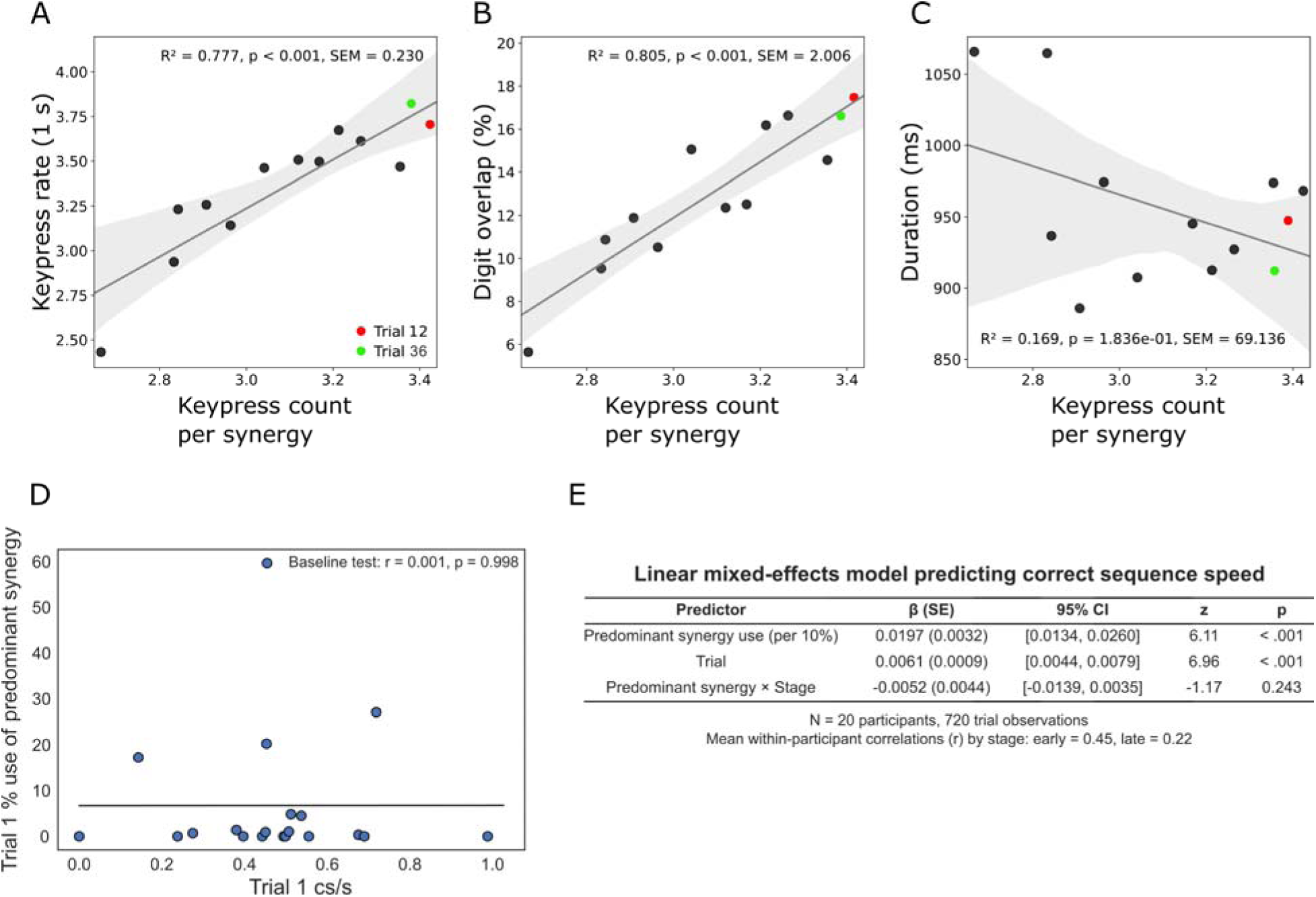
Relationship between the mean number of digits recruited per synergy and corresponding kinematics during early learning (trials 1–12) related to Figure 2. The emergence of higher-order (multi-digit) synergies was associated with progressively faster keypress rates (**A**, R² = 0.777, p < 0.001) and increased temporal overlap of digit movements (**B**, R² = 0.805, p < 0.001), but not with changes in synergy duration (**C**, R² = 0.169, p = 0.184). Data points from Trial 36 (green). We tested whether expression of the predominant synergy simply reflected faster typing. Across participants, Trial 1 typing speed did not predict Trial 1 expression of the predominant synergy (r = 0.0007, p = 0.998; **D**), arguing against the interpretation that expert-like synergy structure is merely a mechanical consequence of baseline speed. **E**. We further assessed the within-participant relationship between synergy use and performance using a linear mixed-effects model with participant as a random effect. Predominant synergy use significantly predicted typing speed above and beyond the effect of trial (β (SE) = 0.0197 (0.0032); 95% CI = 0.0134, 0.0260; z = 6.11; p = <0.001), while trial also showed an independent positive effect (β (SE) = 0.0061 (0.0009); 95% CI = 0.0044, 0.0079; z = 6.96; p = <0.001), consistent with learning across practice. The synergy × stage interaction was not significant (β (SE) = −0.0052 (0.0044); 95% CI = −0.0139, 0.0035; z = −1.17; p = 0.243), indicating that the synergy–performance relationship did not differ reliably between early and late learning. Consistent with this, the correlation between synergy use and typing speed was moderate early in learning (r = 0.45) and weaker later (r = 0.22). Together, these findings indicate that synergy structure captures variance in skill beyond what is explained by skill speed. Mean % accuracy errors across participants was 3.1±0.51% (SEM).

**Figure S3.**
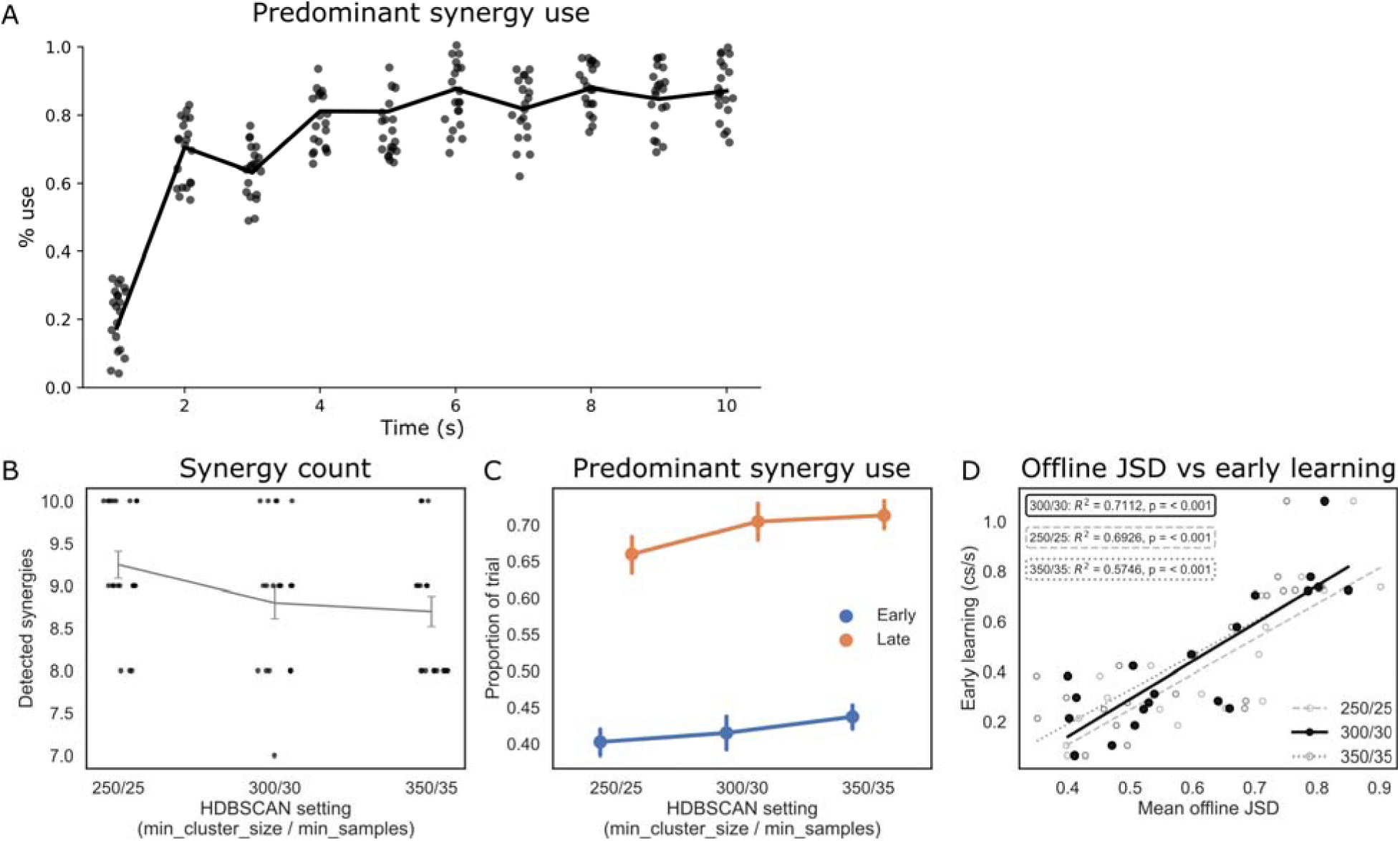
A. No sign of reduction in use of predominant multi-digit synergies late in training, related to Figures 3, 4 and 5. We examined whether accumulating fatigue during the 10-s practice interval of a late trial (Trial 35)—at the end of Day 1, when fatigue would be most likely—reduced the percentage use of the predominant multi-digit synergies (see **Figure. 4**, **Figure. 5**). No such deterioration was observed: multi-digit synergies remained prominently and stably expressed throughout the 10-s interval of this late training trial in all participants. Thus, we found no evidence that accumulating fatigue measurably influenced multi-digit synergy use. **B, C and D**. **Use of nearby sets of UMAP/HDBSCAN hyperparameters rendered comparable results**. **B. Synergy count.** The number of inferred synergies was stable across alternative HDBSCAN parameter settings, indicating that the identified synergy structure was not driven by local changes in clustering thresholds. **C. Predominant synergy use.** The increase in predominant synergy use from early to late learning was preserved across parameter settings, with clear early/late separation in all cases. **D. Offline divergence and performance.** The association between offline JSD and average typing speed was also preserved across parameterizations, with similar effect sizes and significance levels despite minor changes in slope and dispersion. Together, these analyses show that the main findings are robust across a local range of UMAP/HDBSCAN hyperparameters.

**Figure S4.**
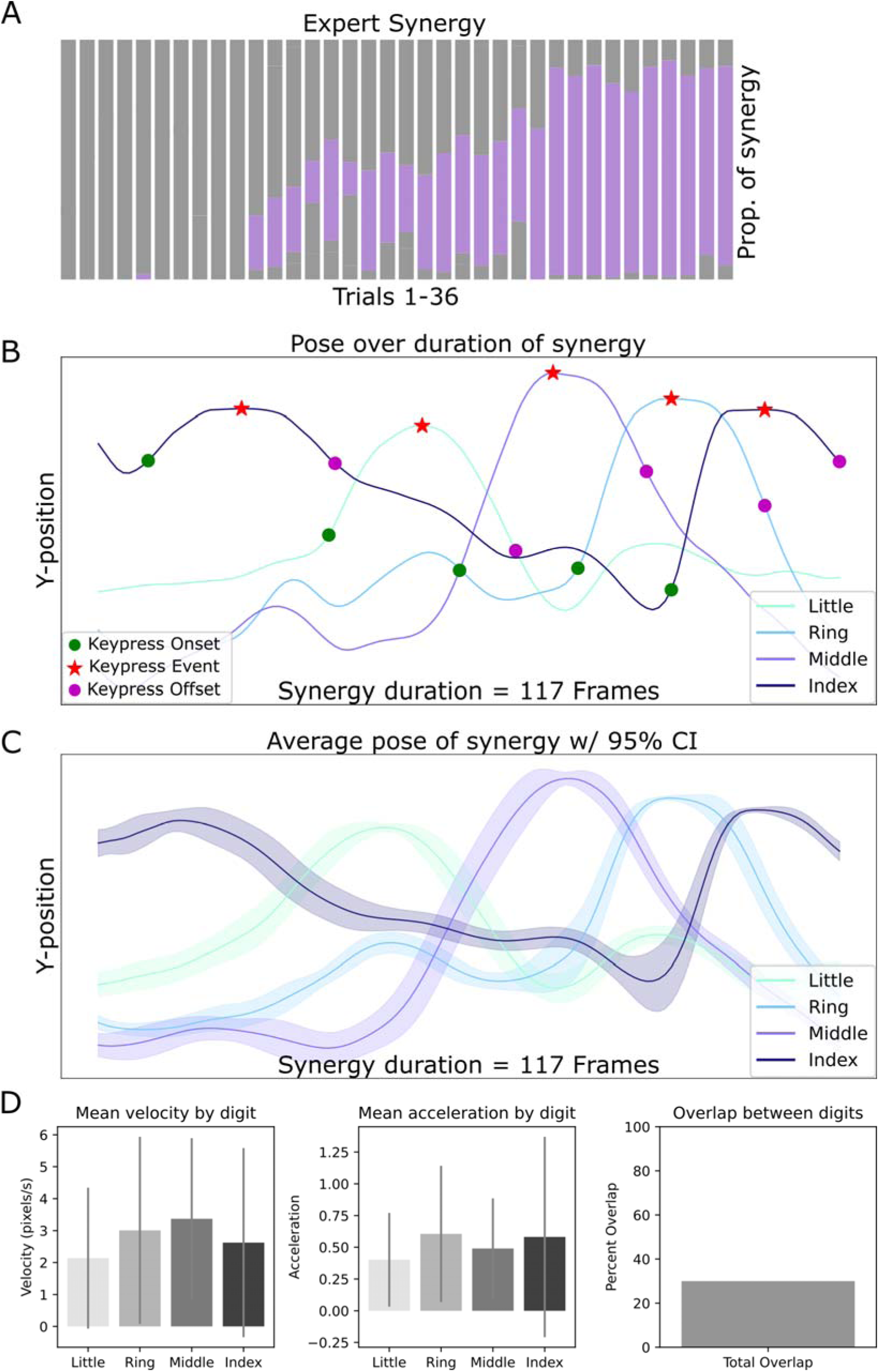
Kinematic Profile of an expert synergy, related to Figure 4 (same subject as shown in Figure 4). **(A)** A single expert synergy across 36 practice trials. Each vertical bar represents one trial, indicating the % use per trial of this synergy as training progressed. This particular synergy encompasses all keypresses in the learned sequence. **(B)** Individual digit kinematics (Y-position, camera frame pixels) during execution of a single 5-keypress (red stars) expert synergy (117 frames, 975ms). **(C)** Mean digit trajectories of the same expert synergy averaged across all occurrences in the course of training (95% confidence intervals). The pattern highlights stable and consistent multi-digit coordination for the expert synergy once it is created. **(D)** Kinematic features of this expert synergy averaged in the course of training (±SDM): mean velocity (pixels/frame, left), acceleration (pixels/frame², middle) and digit overlap (% of frames with digit overlap in the course of training).

**Figure S5.**
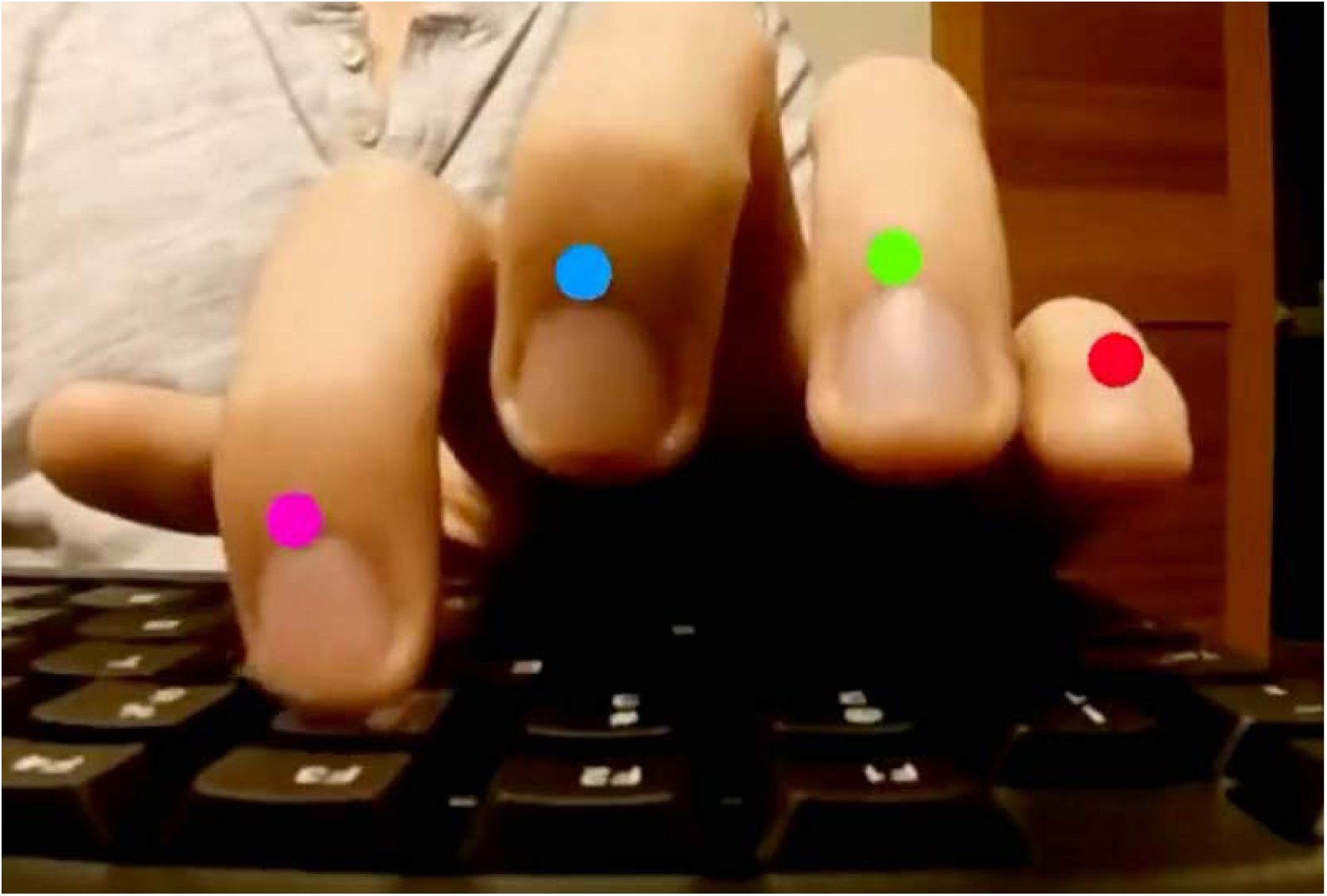
Video with pose-estimated labels at full-speed, related to Figure 4. The video displays the predominant expert synergy illustrated in **Figure S4** (117 frames), repeated consecutively 10 times. This synergy encompasses all keypresses in the learned sequence. Video 1 shows the synergy at its original temporal scale (total duration ≈10 s).

**Figure S6.**
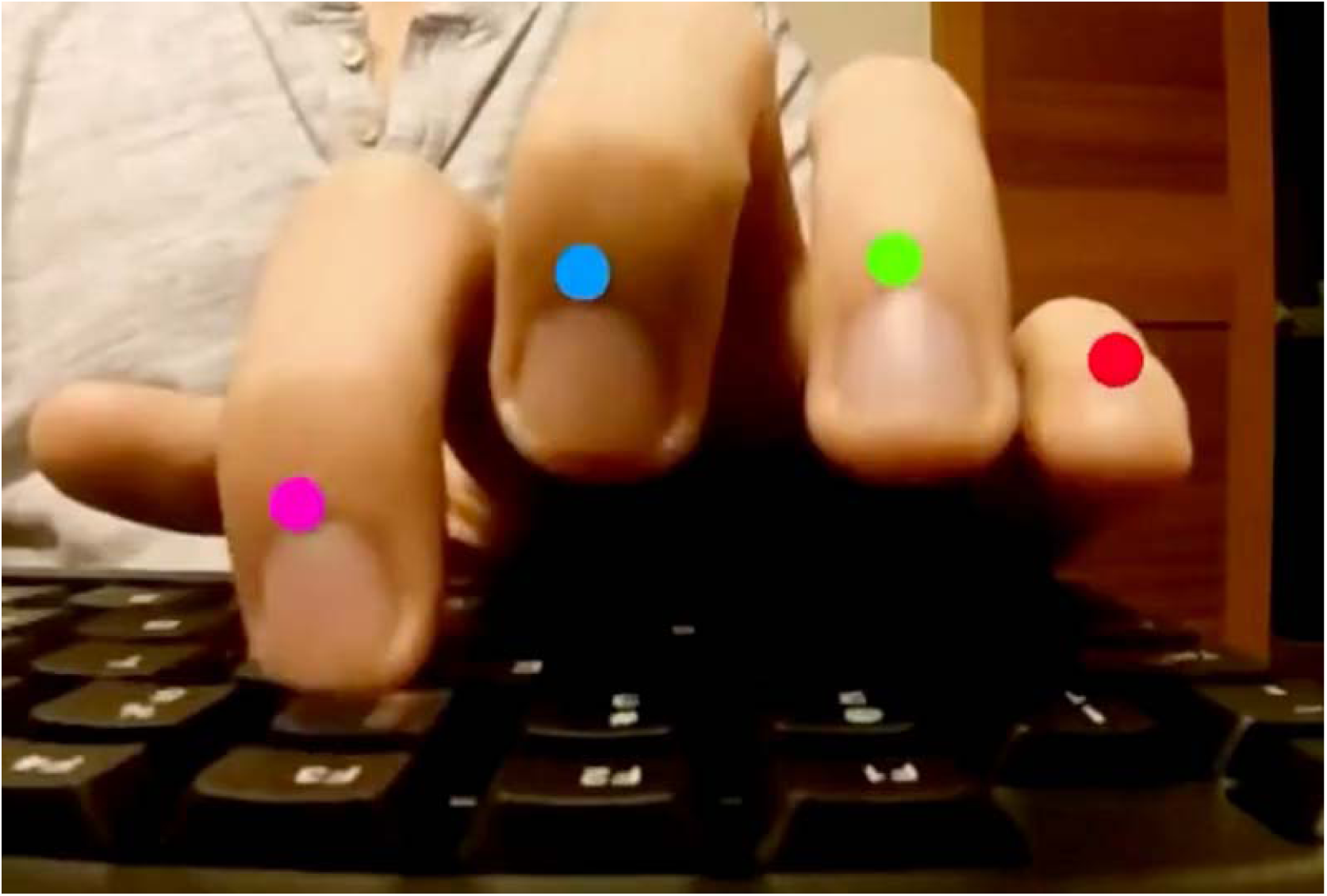
Video with pose-estimated labels at half-speed, related to Figure 4. The video displays the predominant expert synergy illustrated in **Figure S4** (117 frames), repeated consecutively 10 times. This synergy encompasses all keypresses in the learned sequence. This video presents the same synergy as Figure S5 at half-speed (total duration ≈ 20 s), affording clearer visualisation of the underlying temporal dynamics and inter-keypress coordination.

**Figure S7.**
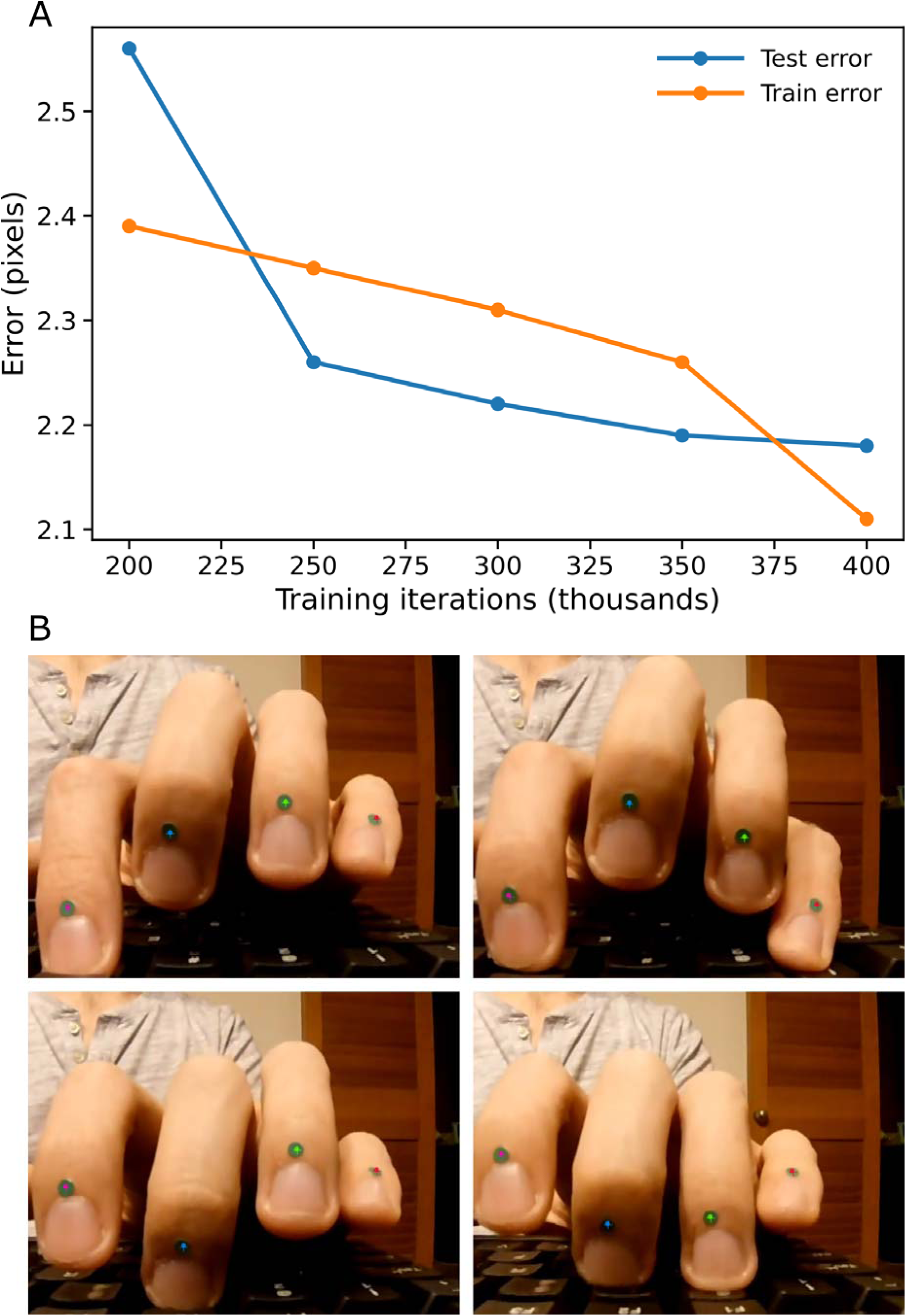
DeepLabCut network training performance and example marker labelling, related to STAR Methods. **(A)** A DeepLabCut neural network was trained to track digit movements using default parameters (400,000 training iterations, RGB input, pairwise terms enabled, without additional supervision). Network performance was evaluated using 1 shuffle split. Root Mean Square Error (RMSE) decreased steadily with training iterations, reaching 2.12 pixels on the training set and 2.18 pixels on the held-out test set (image resolution: 1280 × 720). Error was assessed at intervals of 25,000 iterations. The stability and convergence of both training and test errors indicate that the algorithm learned successfully. (**B**) Representative video frames showing predicted marker placements (coloured dots) on the fingernails of the index, middle, ring, and little fingers. Each marker corresponds to a tracked key anatomical point manually defined during training. Marker predictions shown here were generated by the trained DeepLabCut model and used as input for downstream kinematic analyses of digit position and motion.

During the preparation of this work the authors used GPT-5.2 for editing purposes. After using this tool/service, the authors reviewed and edited the content as needed and take full responsibility for the content of the published article.

